# A data-fusion approach to identifying developmental dyslexia from multi-omics datasets

**DOI:** 10.1101/2023.02.27.530280

**Authors:** Jackson Carrion, Rohit Nandakumar, Xiaojian Shi, Haiwei Gu, Yookyung Kim, Wendy H. Raskind, Beate Peter, Valentin Dinu

## Abstract

This exploratory study tested and validated the use of data fusion and machine learning techniques to probe high-throughput omics and clinical data with a goal of exploring the etiology of developmental dyslexia. Developmental dyslexia is the leading learning disability in school aged children affecting roughly 5-10% of the US population. The complex biological and neurological phenotype of this life altering disability complicates its diagnosis. Phenome, exome, and metabolome data was collected allowing us to fully explore this system from a behavioral, cellular, and molecular point of view. This study provides a proof of concept showing that data fusion and ensemble learning techniques can outperform traditional machine learning techniques when provided small and complex multi-omics and clinical datasets. Heterogenous stacking classifiers consisting of single-omic experts/models achieved an accuracy of 86%, F1 score of 0.89, and AUC value of 0.83. Ensemble methods also provided a ranked list of important features that suggests exome single nucleotide polymorphisms found in the thalamus and cerebellum could be potential biomarkers for developmental dyslexia and heavily influenced the classification of DD within our machine learning models.

## Introduction

### Molecular and Analytical Advances. Data Fusion

High-throughput technologies are continuously advancing, enabling researchers across various fields to utilize genomics, transcriptomics, and metabolomics for instance, to explore complex diseases and disabilities from a comprehensive perspective (1–21). When clinical data is not sufficient or practical enough to make a diagnosis, it is important to have the sufficient methodology and knowledge to utilize these multi-omics tools in making better informed decisions. Viewing complex neurodevelopmental disorders such as developmental dyslexia (DD), autism spectrum disorder (ASD), attention deficit hyperactivity disorder (ADHD), and Alzheimer’s disease through this multi-omics perspective can provide a much deeper understanding of not only the etiology of the disease or disorder but also its pathogenesis and pathophysiology (1, 2). Machine learning (ML) and deep learning (DL) techniques can be very effective at processing single domain datasets, but these high-dimensional and heterogenous datasets seen in multi-omics research can be very complex and noisy for traditional ML methods (4). To address these challenges, data fusion techniques have begun to provide insight on systems biology research by efficiently representing multi-omics data while also extracting the key features.

Yuan et al. (22) tested and validated various data fusion models to assist in the prognosis and clustering of various cancers utilizing not only traditional clinical data but also molecular data like DNA methylation, mRNA, microRNA, and protein expression. Minoura et al. (23) developed a mixture-of-experts model to extract biologically meaningful variables and predict surface protein expression for various cells using single-cell transcriptome, epigenome, and proteome data. Using new deep learning methodologies to explore systems biology data is still new but has shown potential in many disorders (4, 22, 23).

The two most common data fusion techniques are data-level and decision-level fusion (24). Early fusion (data-level fusion) is designed to represent the relationship between modalities in a meaningful way and joins the data while keeping the semantics of each dataset (24). When dealing with heterogeneous datasets, the links between the core data contain fundamental information that can be used to deep dive into the system being explored (24). In patient specific data like clinical and bioinformatics data, these hidden links between different data types can provide researchers the insights into the entire system or pathway instead of specific time points (25). The second form of data fusion is known as late fusion, (decision-level fusion) (2, 24–26). This can be best represented by mixture-of-experts (MoE) models, voting classifiers, stacking classifiers, deep neural networks, etc. (26). Obtaining a consensus from heterogenous classifiers provides a more robust process by leveraging the strengths of the models while also mitigating their weaknesses. Assigning various models to respective modalities can improve generalization due to the division of labor among experts/models.

### Developmental Dyslexia

Developmental dyslexia (DD) is a specific reading disorder that is characterized by difficulties in word recognition, spelling, and phonological processing (27). Dyslexia can be categorized into two main branches, developmental and acquired dyslexia. Reading difficulty resulting from a brain injury is referred to as acquired dyslexia (28). Developmental dyslexia can have genetic components or arise from environmental causes before or during birth (29). The prevalence of developmental dyslexia ranges from 5-10% in school aged children, depending on diagnostic criteria, making this the most common learning disability in US school children (29–31). In various studies, several traits have been reported as comorbid with dyslexia, such as attention deficit hyperactivity disorder (ADHD) (32–35), diminished general processing speed (36–38), and slowed performance during tests of rapid naming (39).

It has been estimated that the sibling recurrence risk of DD generally is 43-60%, while the sibling recurrence risk of DD when both parents have been diagnosed with DD is 77% (31, 40). This suggests a heritable component of DD. Previous studies have nominated potential susceptibility genes, of which four are consistently being reported, *DNAAF4, KIAA0319,DCDC2*, and *ROBO1* (31, 40, 41). *ROBO1* and *DCDC2* have direct involvement in many neurodevelopmental processes like axon growth and neuronal migration, and they may even have a role in the proper development of the corpus callosum, while *KIAA0319* is also assumed to play a direct role in developmental processes like cell adhesion (41, 42). Although some of these genes are not fully understood on a biochemical level, they are all highly expressed in the brain (41, 42). Various structural MRI studies on participants with and without DD have also shown significant differences between the two groups in areas of the brain like the bilateral insular cortex, cerebellum, cerebral cortex, and the thalamus (43–46). Functional MRI studies have also shown that areas like the thalamus respond abnormally in dyslexic individuals, with signs of reduced connectivity from regions like the left visual thalamus to specific cerebral cortex regions in dyslexic individuals (44, 45).

It is also thought the thalamus plays a role in the neural noise hypothesis, that individuals with dyslexia are more susceptible to neural hyperexcitability, leading to difficulty filtering out irrelevant stimuli (33). Since the thalamus plays a role in filtering out irrelevant information supporting various gating process in the brain, it is possible that individuals with functional disruptions in the thalamus could experience heighted “noise” and difficulty with extracting the essential visual and phonological information that is needed for learning to associate written word forms with word meanings (33, 47). Another DD hypothesis is the cerebellar deficit hypothesis, which suggest individuals with DD may have difficulties reading due to developmental abnormalities of cerebellar function (48, 49). The cerebellum supports cognitive functions like the brain’s timing mechanism that helps coordinate the flow of information between different brain segments (48). It has been observed that multiple cerebellar areas play a role in reading and can even provide enough insight to make predictions on an individual’s reading abilities (50). DD is a very complex learning disability that incorporates developmental issues in various parts of the brain. Finding novel genes highly expressed in these regions of the brain may give researchers more insight and new targets to explore.

In this study, we developed and validated various ML/data-fusion models at the data and decision level to find the best way to incorporate heterogenous multi-omics datasets. Classifying dyslexic and non-dyslexic individuals can be difficult due to the complexity of the disorder, and with only clinical diagnostic approaches currently available, this exploratory study aims to prove a bioinformatics data-fusion approach can also provide accurate results. With ML techniques identifying the most influential features, feature importance was used to validate the biological importance of the most influential features.

## Methods

### Participants and Phenotypic Assessments

This project was completed with the oversight and approval of the Institutional Review Boards at the University of Washington and Arizona State University as previously mentioned (51, 52). Adults gave written consent and parents gave written permission for their minor children to participate. Altogether, 89 participants were included in the study, 44 males and 45 females, with an average age of 32.45 and a standard deviation of 19.74. Thirty-six participants were in, 2- or 3-generational families with dyslexia, with the average family size being 5.14 ranging from 4 to 7 (53).

To be included in the dyslexia group, participants were required to score below −1 SD in at least one of five measures of reading and/or spelling, described below. To be included in the typical control group, participants were required to score above −1 SD in all five measures. All participants were required to be free from any developmental or sensory condition that could introduce confounding.

Sight word recognition and nonword decoding were assessed under untimed conditions with the Word Identification (WID) and Word Attack (WATT) subtests of the Woodcock Johnson Reading Mastery Test, Third Edition (54) and under timed conditions using the Sight Word Efficiency (SWE) and Phonemic Decoding Efficiency (PDE) subtests of the Test of Word Reading Efficiency (TOWRE) (55). Spelling ability was assessed with the Spelling subtest of the Wechsler Individual Achievement Test – Second Edition (WIAT-2) (56).

In addition to assessments of reading and spelling, participants were asked to produce rapid sequences of syllables (“papapa …,” “tatata …,” “kakaka …”) to measure motor speeds in a motor speech task. Average syllable durations were computed and transformed into z scores based on published norms (57) To measure rapid naming speeds, the RAN/RAS: Rapid Automatized Naming and Rapid Alternating Stimulus Test was administered (58). Participants named the items in an array of 50 as rapidly as possible. This test includes the following subtests: Letters, Numbers, Colors, Objects, Letters alternating with Numbers, and Letters alternating with Numbers and Colors. For each subtest, completion time is converted to a standard score based on participant age.

### Phenotypic/Clinical Data

Due to ranges in age and other demographic features, only the standardized scores of each clinical test mentioned above were used in the analysis and the raw scores were ignored. Any participant or feature with more than 20% missing values was removed from the analysis. As mentioned above, the resulting missing values were imputed with the average score for the respective phenotypic trait. These results were then scaled using the MinMaxScaler from Sklearn to transform the values to a range between 0 and 1 while keeping the original shape and distribution. This resulted in 58 participants with 47 phenotypic/clinical measurements, of whom 25 were unaffected and 33 were dyslexic.

### Genomics Data

Participants provided blood samples and exome sequencing was completed at the University of Washington Center for Mendelian Genomics (UW CMG) using the Illumina HiSeq 4000 sequencer and the CMG processing pipeline. The CMG processing pipeline consists of RTA (v2.7.6) for base calls generated in real-time, unaligned BAM files produced by Picard ExtractIlluminaBarcodes and IlluminaBasecallsToSam, and finally BWA (Burrows-Wheeler Aligner; v0.7.10) to assist in aligning BAM files to the human reference hg19hs37d5.

Read-pairs mapping within ± 2 standard deviations of the average library size (~150 ± 15 bp) were included while any read-pairs outside of this range (135 bp ~ 165 bp) were removed from further analysis. Picard MarkDuplicates (v1.111) was then used on all aligned data to remove duplicate reads that happen to have similar start positions. To adjust for indels and poor alignment around indels, GATK IndelRealigner (v3.2.2) was used for indel realignment. Finally, GATK BaseRecalibrator (v3.2.2) was used to recalibrate base qualities.

GATK HaplotypeCaller (v3.7) was used for variant detection and genotyping, resulting in the variant data containing genotype data for each individual sample. Further analysis included using the GATK VariantFiltration (v3.7) tool to mark sites of lower quality/confidence when aligned to the hg19 reference genome. Variants were annotated and allele frequency was reported using the (VEP) (59) and Combined Annotation Dependent Depletion (CADD) (60) respectively.

After sequencing the exomes of the participants and compiling a VCF file, an allele frequency PLINK (61) analysis was conducted and only significant (p< 0.02) single nucleotide polymorphisms (SNPs) were collected for future analysis. The P-value cutoff of 0.02 was used to select the 332 top scoring SNPs for further analysis. One-hot encoding was then used to better represent the participants DNA where 0, 1, and 2 represent homozygous for the reference allele, heterozygous, and homozygous for the alternative allele respectively. Of the 89 individuals that participated in this study, 54 had quality sequencing data sufficient for this study and 332 distinct SNPs were analyzed for every participant. Five of the qualified individuals were reported as ‘undetermined’ regarding their dyslexia diagnosis so they were removed from the analysis. The remaining 49 participants included 29 with dyslexia and 20 without dyslexia. *Metabolomics Data*

Participants provided saliva samples (2 ml), which were collected and stored at −80°C. Frozen saliva samples were thawed overnight at 4°C and 100 μL of each sample was used for further analysis. To isolate proteins and metabolites from the saliva, 500 μL of methanol and 50 μL standard solution (1,810.5 μM ^13^C_3_-lactate and 142 μM ^13^C_5_-glutamic acid in PBS; Sigma-Aldrich) were added to each sample. The mixture was vortexed (10 sec), stored at −20°C for 30 min, and was followed by centrifugation (14,000 RPM, 10 min, 4° C). A CentriVap Concentrator (Labconco) was used to transfer and dry the supernatants. The dried samples were then restored in a fixed volume (150 μL) of reconstitution buffer (40% (v/v) PBS (GE Healthcare), 60% (v/v) acetonitrile (Fisher Scientific)).

All LC-MS/MS procedures were performed on an Agilent 1290 UPLC-6490 QQQ-MS (Santa Clara) system. Both negative and positive ionization modes were used and required 10 μL and 4 μL respectively for analysis. Liquid chromatography purification was performed using hydrophilic interaction chromatography with a Waters XBridge BEH Amide column (Waters Corporation). A fixed flow rate (0.3 mL/min) was selected, auto-sampler temperature was kept at 4 °C, and the column was set at 40 °C. The mobile phase was composed of Solvents A (10 mM ammonium acetate, 10 mM ammonium hydroxide in 95% H_2_O/ 5% acetonitrile) and B (10 mM ammonium acetate, 10 mM ammonium hydroxide in 95% acetonitrile/ 5% H_2_O). After the initial isocratic elution with 90% B, the UPLC decreased solvent B to 40%, held the gradient for 4 minutes, and then return to 90%. This process was repeated for the following samples. Mass spectrometry was performed using an electrospray ionization source while multiple-reaction-monitoring (MRM) mode was used for more sensitive data collection. Agilent Masshunter Workstation (Santa Clara, CA) software was used to control the LC-MS system and extract MRM peaks. Statistical analysis of the metabolites was performed using MetaboAnalyst (62) and the metabolite levels were transformed using a log10 transformation. Linking these metabolite levels to the respective participant and class label were conducted using Pandas software in Python (v3.8.10).

Of these 296 metabolites, 28 contained over 20% missing values and were subsequently removed from the study. For any remaining participants with missing specific metabolite levels, the missing values were imputed with the average metabolite level according to the other individuals in the study. These results were then scaled using the MinMaxScaler from Sklearn to transform the values to a range between 0 and 1 while keeping the original shape and distribution. From the original study, 26 of the 89 participants, 9 unaffected and 17 with dyslexia, had sufficient metabolomic data collected.

### Joint Data

Two datasets were then created by joining exome data with metabolite data and another dataset that contains the exome, metabolomic, and phenotypic datasets. Only a small subset of participants had quality data in both datasets, and this resulted in a small sample size (n = 19) in both datasets. Of the 19 participants in the analysis, only 4 were unaffected and 15 were individuals with dyslexia. Due to the imbalanced data, SMOTE (63) was used to replicate and create new data points for the minority class (random state 18 with k-nearest neighbors = 3). After utilizing SMOTE, the new balanced datasets included 30 instances where 15 were unaffected and the other 15 had dyslexia. These 30 participants provided 671 attributes including exome, metabolomic, and phenotypic features.

### Data-level fusion (Figure 1a)

Both multi-omics datasets generated by SMOTE, Exome-Metabolite and Exome-Metabolite-Phenotypic, were split 80/20 for a training and test dataset respectively and analyzed using various machine learning techniques including neural network (multi-layer perceptron, MLP), random forest (RF), K-nearest neighbor (KNN), and ensemble learning techniques including AdaBoost (ADA). To optimize the various models for each dataset, grid search was implemented to tune the model and various parameters with respect to ROC AUC and paired with 10-fold cross validation. The grid search would return the most optimal model for that respective classifier. The testing dataset was then used as input for the resulting optimized models to give more accurate metrics like F1 score, accuracy, AUC. Interpretable tree-based models like AdaBoost and random forest were further examined, and features were ranked and plotted to visualize the importance of each feature. Feature importance is calculated with Gini importance using Sci-kit Learn in Python, allowing the impurity of nodes/features to be considered and reward/increase the importance of features that have purer splits. The feature importance algorithm is detailed in the Appendix. The minor allele frequencies for the important exome SNPs were determined via GnomAD v2.1 (64)

**Figure 1.**
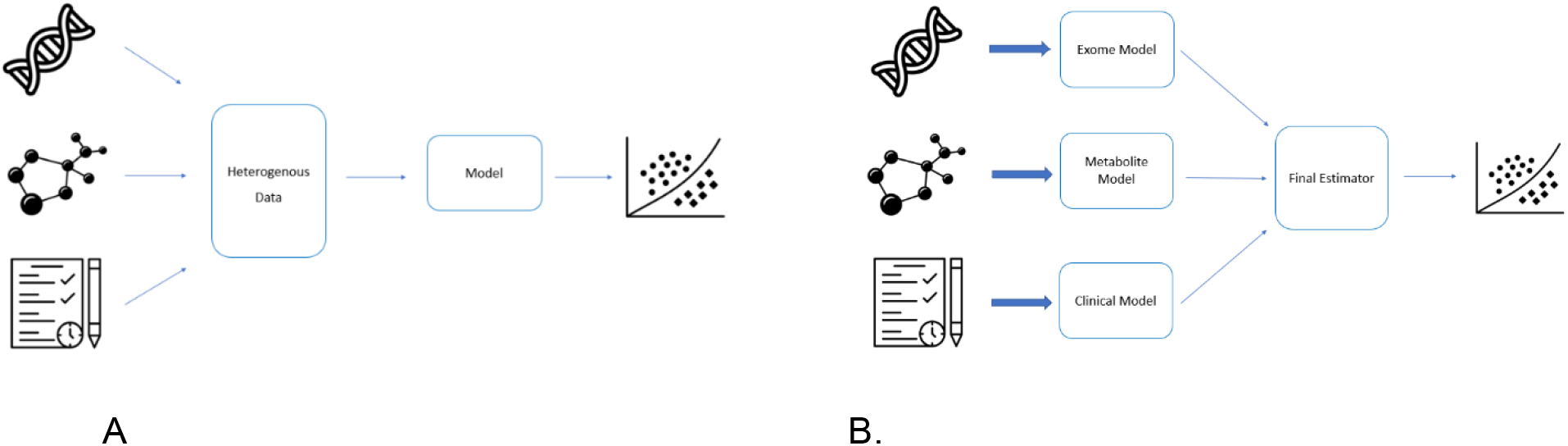
A. Visual representation of data-level fusion. B. Visual representation of decision-level fusion.

### Decision-level fusion (Figure 1b)

Several machine learning models were trained on the single-omic data sets individually. MLP, RF, KNN, ADA, and gaussian naive bayes (NB) models were trained on 80% of their respective single-omic dataset and the remaining 20% of samples were used for a testing set as mentioned before. Each of these models listed above were tested thoroughly with grid search to find the most optimal parameters for their respective dataset. A decision level classification model was then implemented by stacking the most optimal single-omic models and classified by a final predictor. The best classification model for exome data, metabolome data, and phenome data were implemented in the stacking classifier based upon F1 score and the final estimator was tested between Logistic Regression and Gaussian Naive Bayes. These ensemble models were then trained on 80% of the joint multi-omics datasets and tested with the other 20% as mentioned above.

## Results

### Data-level fusion

Table 1 summarizes the performance results of the top preforming models listed in the methods for their respective dataset. When looking at early data-fusion for exome and metabolome data, ensemble techniques like AdaBoost (learning rate of 0.0005 and 25 estimators/stumps) paired with decision tree (max depth of 4 and max leaves of 4) resulted in a model that had 86% accuracy, an F1 score of 0.86, and an AUC value of 0.92 (Figure 2). Other models like deep neural networks (3 hidden layers with 300, 100, and 10 nodes in each layer) were also able to achieve 86% accuracy with an F1 score of 0.89. Random forest and KNN performed the same achieving 71% accuracy, an F1 score of 0.8, and an AUC value of 0.67.

**Table 1.** Performance metrics for the most optimal model according to GridSearchCV

| Data-level | MLP<br>(300,100,10)<br>Alpha = 0.001 | KNN<br>K = 3 | RF<br>Max Depth = 4<br>Max Leaves = 4 | ADA<br>Estimators = 25<br>Alpha = 0.0005 |
| --- | --- | --- | --- | --- |
| Exome+Metabolome | Accuracy: 0.86<br>F1: 0.89<br>ROC AUC: 0.83 | Accuracy: 0.71<br>F1: 0.80<br>ROC AUC: 0.67 | Accuracy: 0.71<br>F1: 0.80<br>ROC AUC: 0.67 | Accuracy: 0.86<br>F1: 0.86<br>ROC AUC: 0.92 |

| Data-level | MLP<br>(10, 25, 10) | KNN<br>K = 3 | ADA<br>Estimators = 7<br>Alpha = 0.001 |
| --- | --- | --- | --- |
| Exome+Metabolome<br>+Phenome | Accuracy: 0.83<br>F1: 0.89<br>ROC AUC: 0.75 | Accuracy: 0.83<br>F1: 0.89<br>ROC AUC: 0.88 | Accuracy: 0.83<br>F1: 0.67<br>ROC AUC: 0.75 |

**Figure 2.**
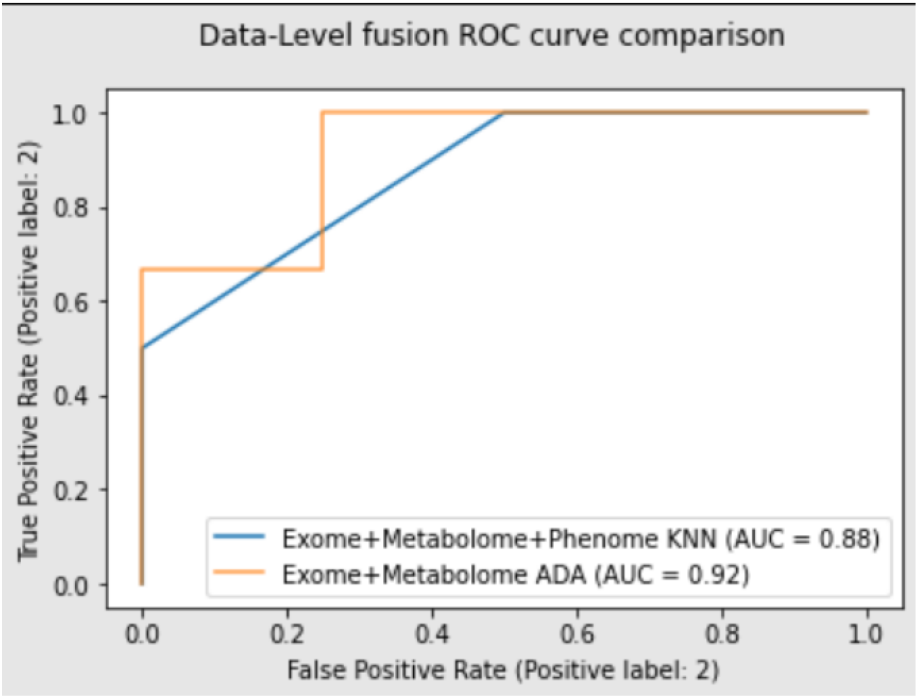
The ROC curve comparing the various early fusion models. This ROC curve can show the effects of adding phenotypic/clinical data into the model.

When performing classification on the exome + metabolome + phenome data, MLP and KNN were the most optimal models. A MLP model with 3 hidden layers consisting of 10, 25, and 10 nodes in each layer was able to achieve 83% accuracy, a F1 score of 0.89, and an AUC value of 0.83. KNN models with K equal to 3 resulted in matching accuracy and F1 score, 83% and 0.89, but had an AUC value of 0.88 (Figure 2). AdaBoost (learning rate of 0.001 and 7 estimators/stumps) was able to achieve 83% accuracy and an AUC value of 0.75 but had a F1 score of 0.67.

Figure 3 depicts feature importance in the interpretable AdaBoost models for the respective datasets. When exploring exome and metabolome merged data, Deoxyuridine monophosphate (dUMP) and the variant rs10988589 were identified as the most important. When using hg19 for alignment of the SNPs, rs10988589 is located at chr9:130092203 in the *GPR107* gene and has a minor allele frequency of 0.227. rs62128466 (a SNP found at chr19:1487562 in the *PCSK4* gene with a minor allele frequency of 0.00003) was the third most important feature, while metabolites lenalidomide and pyridoxine were fourth and fifth (equally) most important features. 6-Phosphogluconic acid, rs10215048 (a SNP found at chr7:150793348 in the *TMEM176B* gene with a minor allele frequency of 0.137), acetylglucosamine acid, muconic acid, and cystenine also showed minor importance when classifying individuals based soley on exome and metabolome data.

**Figure 3.**
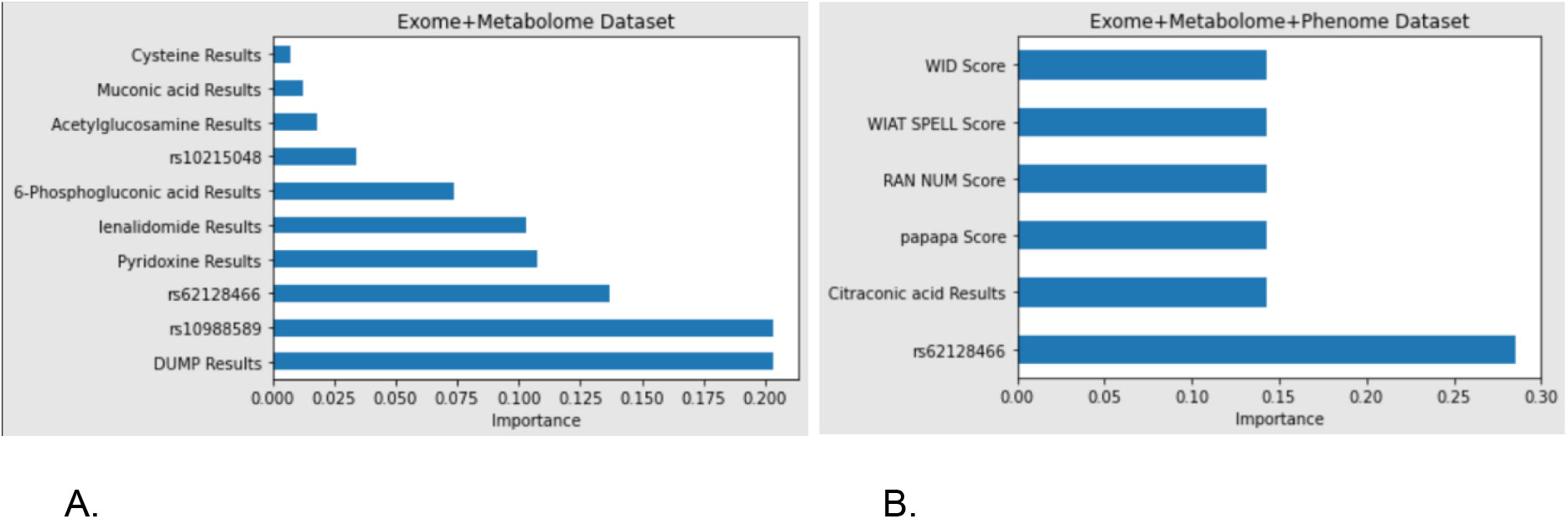
Feature importance of the top ranked attributes. A. Exome and metabolome joint dataset. B. Exome, metabolome, and phenome joint dataset

For the dataset containing phenotypic data, the top six features were also plotted and ranked based on importance. The most important feature was rs62128466 (a SNP found at chr19:1487562 in the *PCSK4* gene with a minor allele frequency of 0.00003), followed by 5 features that are all equally important. Citraconic acid, the standardized score of a syllable repetition speed test (papapa), the standardized score of a rapid automatic naming of numbers test (RAN), the standardized score of the Wechsler Individual Achievement Spelling Test (WIAT), and the standardized score of the Word Identification (WID) subtest of the Woodcock Reading Mastery Test - 3.

Figure 4b shows brain-wide gene expression levels for *PCSK4*, the gene that contains the important SNP rs62128466 seen in both feature importance graphs, using the Allen Human Brain Atlas and Neurosynth. Neurosynth shows various gene expression levels in the brain including an increased gene expression in areas correlating to the thalamus and this can be confirmed by the standard MRI images as well in Figure 3a. The other figures, 3c and 3d, also show brain-wide gene expression levels for *TMEM176B* and *GPR107*, which included important SNPs in the genome+exome analysis.

**Figure 4.**
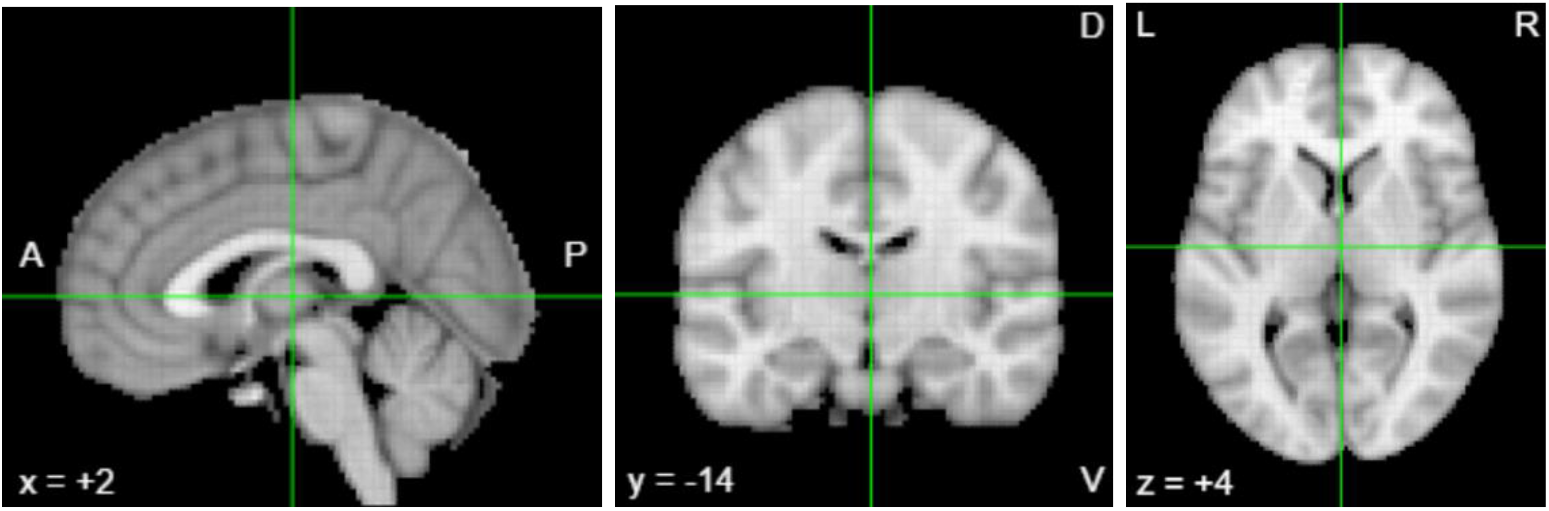
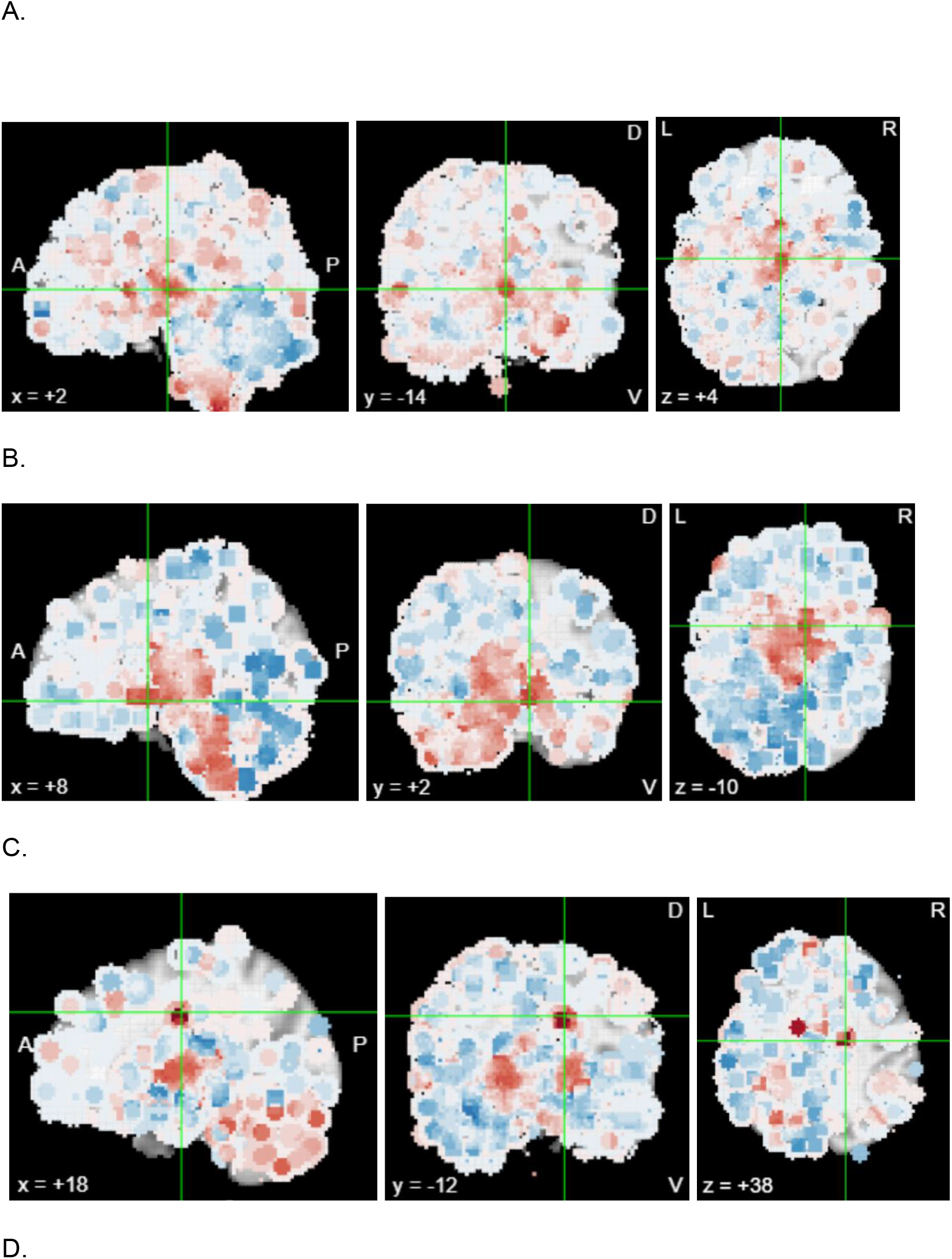
Brain images illustrating the expression pattern of genes with top scoring genomic variants associated with dyslexia. A. Neurosynth MRI image. At location (2, −14, 4) the highest correlated region of the brain is the thalamus. B. Neurosynth *PCSK4* gene expression intensity plotted over MRI image. At location (2, −14, 4) the intensity score is 2.0. C. Neurosynth *TMEM176B* gene expression intensity plotted over MRI image. At location (8, 2, −10) the intensity score is 2.09 and most correlated with the nucleus accumbens. D. Neurosynth *GPR107* gene expression intensity plotted over MRI image. At location (18, −12, 38) the intensity score is 2.52 and most correlated with the cerebellum.

**Figure 4.**
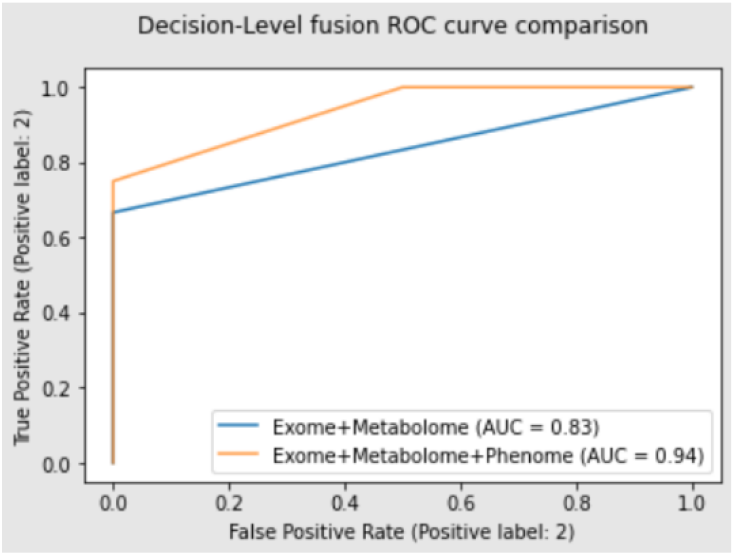
ROC curve comparing the two decision-level fusion models showing the effects of adding the additional datatype (phenome).

### Decision-level fusion

Table 2 summarizes the results from various machine learning techniques on the respective single-omic datasets. Neural networks/multi-layered perceptron (MLP), random forest (RF), K-nearest neighbor (KNN), and AdaBoost (ADA), and gaussian naive bayes (NB) were all trained and tested on separate data subsets. ROC AUC values ranged from 0.5 to 0.75 for exome datasets, 0.4 to 0.7 for metabolite datasets, and 0.75 to 0.97 for phenotypic datasets. For each dataset, (exome, metabolite, and phenotypic) the following models were evaluated and optimized. MLP models optimized with gridsearch returned a model that has 3 different hidden layers with 10 nodes in each hidden level for exome data and a model that has 2 hidden layers with 10 nodes in each for metabolomic and phenotypic data. Optimized random forest models on the single-omic data resulted in trees with a maxDepth of 3 and 25 estimators. K-nearest neighbor algorithms where n = 3 were the most accurate models for the single-omic datasets. The most optimal gaussian naive bayes returned a model with smoothing parameters of 1e-8. ROC curves for each dataset can be seen in Figure 5 and the accuracy, F1 score, and AUC for each model can be found in Table 2.

**Table 2:** Performance metrics for the best model for each respective dataset.

|  | MLP | RF | KNN | GNB |
| --- | --- | --- | --- | --- |
| Exome | Accuracy: 0.75<br>F1: 0.77<br>ROC AUC: 0.75 | Accuracy: 0.75<br>F1: 0.77<br>ROC AUC: 0.75 | Accuracy: 0.75<br>F1: 0.77<br>ROC AUC: 0.75 | Accuracy: 0.50<br>F1: 0.29<br>ROC AUC: 0.50 |
| Metabolome | Accuracy: 0.66<br>F1: 0.50<br>ROC AUC: 0.62 | Accuracy: 0.75<br>F1: 0.77<br>ROC AUC: 0.75 | Accuracy: 0.50<br>F1: 0.0<br>ROC AUC: 0.38 | Accuracy: 0.50<br>F1: 0.0<br>ROC AUC: 0.38 |
| Phenome | Accuracy: 0.83<br>F1: 0.83<br>ROC AUC: 0.94 | Accuracy: 0.92<br>F1: 0.86<br>ROC AUC: 0.91 | Accuracy: 0.92<br>F1: 0.86<br>ROC AUC: 0.97 | Accuracy: 0.83<br>F1: 0.67<br>ROC AUC: 0.75 |

**Figure 5.**
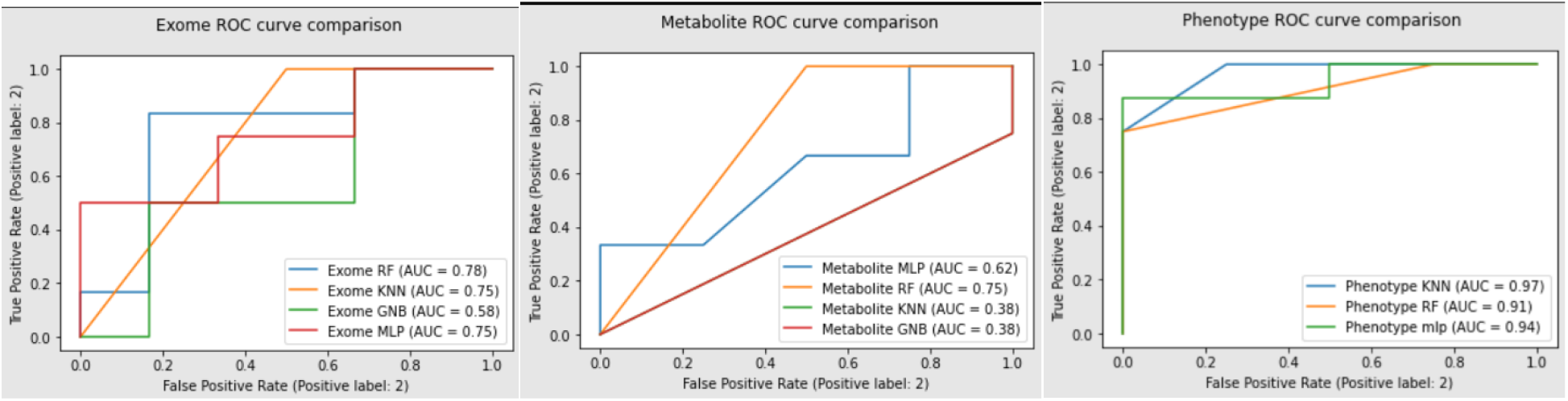
ROC curves comparing the various models for each dataset. A. Exome dataset, B. Metabolite dataset, C. Phenotype dataset.

The best predictors for each individual datatype, based on F1 score, were then used to construct a stacking classifier which resulted in a model that can achieve 86% accuracy with a F1 score of 0.89 for Exome-Metabolite datasets and 92% accuracy with an F1 score of 0.89 for Exome-Metabolite-Phenotypic datasets. Performance results for the modes mentioned above can be found in Table 3.

**Table 3.**
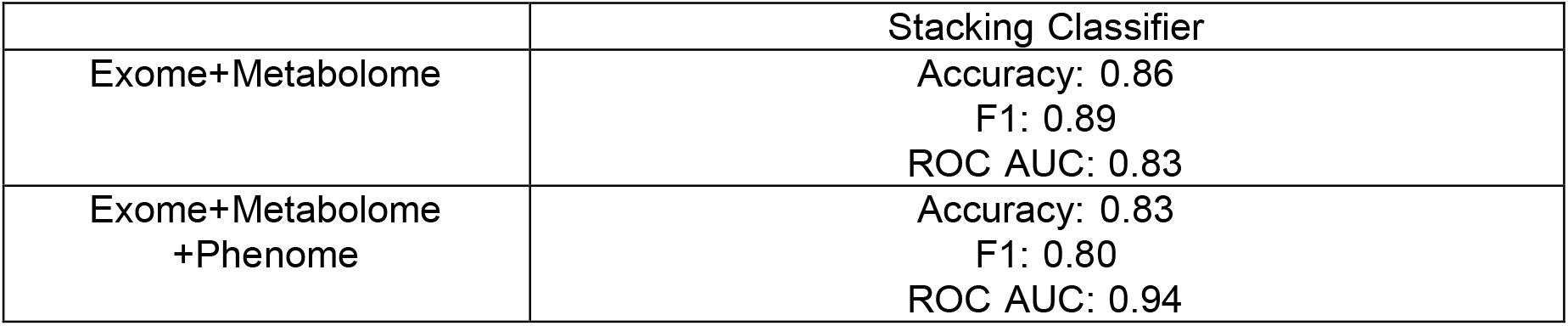
Performance metrics for the decision-level stacking classifiers

## Discussion

In this study, we aimed to create and compare ML models and data-fusion models, at the data (early) and decision (late) level, for their effectiveness in performing disorder classification using a combination of -omics (genomics and metabolomics) and clinical features. We applied these approaches in the context of developmental dyslexia, a disorder believed to have a complex etiology involving multiple genetic and environmental factors. We also explored the use of explainable AI techniques to identify the most influential features in making accurate predictions.

When exploring the single-omic datasets, for exome and metabolome data, we were able to consistently achieve ~75% accuracy using various traditional machine learning models. The phenome/clinical models achieved accuracy upwards of 92% accuracy, which is most probably due to the ground truth diagnosis of DD being based on the individual’s performance on the clinical tests. When exploring the combination of these dataset, the biggest difference was the increase in performance seen in exome+metabolome models. Going from 75% accuracy to 86% accuracy is a significant boost in performance when using only exome and metabolome data. Both deep neural networks and ensemble methods like Adaboost provided high performance metrics at the data-level while at the decision-level, a stacking classifier consisting of the exome KNN model, and the metabolome RF model was able to achieve slightly higher performance metrics as seen above.

The biggest discrepancy seen in the results was the surprising decrease in performance in the models containing clinical (phenome) data when using the data-level fusion approach. Due to the clinical models achieving 0.94 – 0.96 AUC values, we assumed the addition of this datatype would have aided the models in making more educated predictions. Early fusion, data-level fusion, models that were trained only on the molecular data, exome + metabolome, received slightly higher F1 score and accuracy performance metrics compared to the heterogenous data containing both molecular and clinical data. At the data-level, we see a 3.5% decrease in accuracy, 0.03 decrease in F1 score, and a 0.04 decrease in ROC AUC values when the models were exposed to the entire dataset containing the participant’s phenome. The ability to predict DD status in an individual with just the exome and metabolome at 86% accuracy could provide more insight on the etiology and development of dyslexia as well as provide new early detection methods for younger and mentally challenged individuals who cannot take these clinical tests.

At the decision-level we see the expected increase in performance after the clinical data were incorporated resulting in the overall best model over both fusion techniques. The best model was a stacking classifier consisting of the exome KNN model, metabolome RF model, and the phenome/clinical RF model with logistic regression model as the final estimator. Although the F1 score is identical, 0.89, the ROC AUC increased from 0.83 to 0.94 and the accuracy changed slightly from 0.86 to 0.83 when comparing the decision-level exome+metabolome model and the decision-level exome+metabolome+phenome model respectively. This mixture-of-experts method performed better than the individual omic models when looking at exome and metabolome data. The initial 75% accuracy that exome and metabolome data can achieve individually is a fair performance but when you combine the decisions of these two classifiers, the accuracy increases to 86%.

With the feature importance mapping we also determined that the mixture-of-experts methods do not simply weigh certain modalities higher than others but encompasses a true mixture of features from all datatypes. We can also see that the model also chose to weigh SNPs in the *GPR107, PCSK4*, and *TMEM176B* genes, which are expressed in various regions of the brain. These genes have not been reported yet as associated with DD, however, their respective region of high gene expression can be seen in areas that are known to affect components in the brain related to DD hypotheses. Affected regions of the brain like the cerebellum, nucleus accumbens, and thalamus can cause drastic effects in the development of one’s ability to effectively read, write, etc.

The rs62128466 SNP from the *PCSK4* gene was also deemed important throughout both datasets. When analyzing the exome+metabolome data, rs62128466 was the third most important feature, and when analyzing the exome+metabolome+phenome data, rs62128466 was the most first important feature. This very rare SNP (minor allele frequency of 0.00003) is found in the *PCSK4* gene, proprotein convertase subtilisn/kexin type 4, which is shown to have a high expression level in the thalamus. The thalamus has been reported to have direct influence in the diagnosis and development of DD, and looking at Figure 3b, we can see the highest levels of *PCSK4* gene expression are around the center of the left and right thalamus. This potential genetic biomarker could play an influential role on the development and overall function of the thalamus, which may play a crucial role in DD according to the neural noise hypothesis.

The most important SNP in the exome+metabolome data is located in the *GPR107* gene which can be seen in Figure 3d. *GPR107* is a gene that codes for the G protein-coupled receptor 107 and not only does this gene show the highest gene expression levels in the brain out of the other high-importance SNPs, but it also has a high correlation to the cerebellum. G protein-coupled receptors, GPCRs, are major factors in cell communication and recognition pathways and they have been implicated in a variety of diseases as well (65). GPCRs are expressed throughout the brain and can be influential in major senses like taste, smell, and, most importantly, vision (65). Studies have shown that GPR107 is specifically involved with retrograde protein transport which is a very important factor in intracellular regulation (66). The ability to use data-fusion techniques as a process to explore biologically important features from multi-omics datasets could play an important role in identifying new biomarkers or drug/therapy targets.

## Conclusion and Future studies

With high throughput technologies and as the use of systems biology knowledge grows, developing and validating new methods to analyze these heterogenous datasets and incorporate a systems biology point of view can provide more insight and make better predictions as seen above. Due to the diagnosis of DD being reliant on various clinical tests, we can see the clinical/phenome models were very accurate, showing a clinical diagnosis is still the gold standard. Although, for children under the age of 6, or individuals with hearing, slight, and other sensory/developmental abnormalities, they cannot confidently be diagnosed via the clinical tests mentioned in this study. This leads to a need for a more bioinformatics approach compared to the traditional clinical diagnosis. Both data-level and decision-level fusion methods provided better results when analyzing exome and metabolome data, compared to the traditional single-omics ML methods. This pure bioinformatics data-fusion approach has not been reported to our knowledge, in the space of DD, and compared to the 92% accuracy we see in the clinical data, the ability to classify individuals with 86% accuracy at both the data and decision level means exome and metabolome data can also be a good option for making accurate predictions. As states begin to require DD screening in young school children, results are showing these teacher provided screenings, that incorporate some of the clinical tests mentioned in this study, are 69-72% accurate (67). The data-fusion method will not compete with a clinician but can perform better than teacher provided screening.

When adding an additional data type into the dataset, clinical tests, decision-level fusion methods were able to achieve the best predictions and highest performance metrics. The mixture-of-experts/ stacking classifiers models consistently outperformed any other decision-fusion model with our given dataset. Although the data-level fusion methods reacted poorly to adding another modality, the decision-level fusion was able to adjust and better its performance when accounting for the new modality. Due to traditional ML techniques resulting in 92% accuracy when trained on clinical data, the use of data-fusion including clinical data is redundant and only provides more noise for the model to filter through. But we did notice how data-level and decision-level fusion methods react differently when supplied with an additional data type, suggesting decision-level fusion can be more robust and accurate when dealing with various heterogenous datasets. This knowledge can be applied to various fields of research that are incorporating multiple modalities from self-driving cars with multiple sensors to systems biology.

This is the first developmental dyslexia multi-omics study conducted with ML to our knowledge and has been able to not only allow for better screening of DD using a bioinformatics approach but also a better understanding from a biological aspect on why and how the ML models reached their decisions. These methods can be applied to an array of disorders and is not limited to just exome, metabolome, and clinical data. Mixture-of-Expert models provide a versatile approach when dealing with multi-omics datasets and other heterogeneous data. Designing and validating novel data-fusion techniques can give researchers in many fields a new way to explore multiple modalities/datatypes simultaneously. Allowing them to not only achieve higher performance metrics but also get a better understanding of the system.

As mentioned in the methods section, our biggest limitation is the fact that the multi-omics datasets are extremely small (n=30) in this study. Future studies should include a larger sample size and should collect data evenly across both dyslexic and unaffected individuals to eliminate the need of creating synthetic data with SMOTE. This will also allow us to further validate these exploratory methods as well as find new patterns and potential biomarkers for DD. Further, different phenotypic profiles of dyslexia, e.g., neural noise vs. cerebellar underpinnings, should be added to the analysis. This could be done by including more modalities like structural and functional MRI data. As mentioned in the background, there are significant differences between a dyslexic and neurotypical brains and adding another data layer for the algorithms to differentiate against could be beneficial. Adding more diverse modalities may allow data-fusion models to gain even more insight and potentially identify DD and other disorders faster and more accurately than providers and physicians given the right data. This could lead to earlier diagnosis, new treatment approaches that are proactive and personalized, and better health, a translation of principles of precision medicine that is already in progress regarding disorders of spoken language (68). Precision medicine is just one of many fields that can benefit greatly due to the advances of data-fusion and other advanced ML techniques.

## Credit authorship contribution statement

**Jackson Carrion:** Conceptualization, Data curation, Formal analysis, Investigation, Methodology, Software, Validation, Visualization, Project administration, Writing - original draft. **Rohit Nandakumar:** Data curation, Software. **Xiaojian Shi:** Data curation. **Haiwei Gu:** Data curation, Metabolome analysis. **Yookyung Kim:** Data curation. **Wendy H. Raskind:** Resources, Funding acquisition, Writing - review & editing. **Beate Peter:** Data curation, Project administration, Writing - review & editing, Validation, Supervision. **Valentin Dinu:** Conceptualization, Project administration, Writing - review & editing, Validation, Supervision.

## Declaration of competing interest

The authors declare that they have no known competing interests.

## Funding Sources

Grant Number: R01HD088431, Arizona State University JumpStart Fund, University of Washington Royalty Research Fund

## Data and code availability

Input data can be found in the supplementary methods and the code can be found at https://github.com/jtcarrion/multiOmics_DataFusion

## Acknowledgements

Many thanks to the participants for their time and effort.

## Appendix

The feature importance algorithm used by Sci-kit learn has two importance aspects and steps. The first is to identify the Gini importance for each of the features, this can be represented as n_j_ = w_j_C_j_ – w_left(j)_C_left(j)_ – w_right(j)_C_right(j)_, where n_j_ is the importance for the j^th^ feature/node, w_j_ is the weighted number of samples in node j, C_j_ is the impurity value at node j, left(j) and right(j) are the children nodes on the left and right of node j respectively. Due to ensemble methods making various trees, measuring the importance of these features throughout all trees will be calculated as 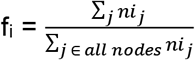, where i is the set of features, j is the current node, and nij is the node importance we measured before. We then can compare f_i_ for all the resulting features and graph the ordered list of features with their respective f_i_ score.

## Notes

https://github.com/jtcarrion/multiOmics_DataFusion

### Competing Interest Statement

The authors have declared no competing interest.

